# Optimal Transport Model for Gas Migration in Pneumatosis Intestinalis

**DOI:** 10.1101/2025.03.10.642225

**Authors:** Arturo Tozzi, Raffaele Minella

**Author notes:** (corresponding author) 1155 Union Circle, #311427 Denton, TX 76203-5017 USA.

## Abstract

Optimal transport models may provide a robust framework for quantifying gas movement from the intestinal lumen into the gut layers in pneumatosis intestinalis (PI), a condition characterized by abnormal gas accumulation within the intestinal wall. We develop a mathematical approach that integrates circular gas diffusion from the lumen with the subsequent dispersion and entrapment of microbubbles in deeper layers. Our diffusion-advection approach establishes a radial concentration gradient that decreases with distance from the lumen. Numerical simulations illustrate the spatial distribution of gas and microbubbles within a representative three-millimeter-thick gut wall, demonstrating a transition from uniform diffusion to localized gas retention. Gas migration initially follows diffusion-driven transport but increasingly becomes influenced by local trapping mechanisms, leading to spatially heterogeneous gas retention. Microbubble formation is modeled as a function of gas entrapment in deeper tissue layers, integrating mechanical effects related to gas concentration and pressure dynamics. Additionally, our model accounts for critical biological and structural factors such as tissue permeability, mucosal integrity, bacterial fermentation, microbial overgrowth and structural heterogeneity, capturing their combined influence on gas retention. Overall, gas migration in PI follows an early-stage circular diffusion pattern, until it encounters structural resistance leading to deep microbubble formation. The density and depth of localized microbubble retention show a statistically significant correlation with gas pressure gradients and permeability variations, highlighting the interplay between transport dynamics and tissue features. Our quantitative representation provides a structured approach for refining diagnostic assessments of PI and improving the evaluation of disease severity based on underlying transport mechanisms.

## INTRODUCTION

Pneumatosis intestinalis (henceforward PI), characterized by the presence of gas within the intestinal wall, is often associated with physiological or pathological conditions including ischemia, infection and mechanical obstruction (Ho et al., 2007; Muchantef et al., 2013; Kaya et al., 2014; Labbé and Hafiani, 2015; Sugihara and Okada, 2017; Suzuki et al., 2017; Maulik et al., 2019; Castellana et al., 2023). Current imaging techniques such as computed tomography and ultrasound allow for the visualization of intramural gas but lack the ability to quantitatively assess its spatial distribution and progression (Heng et al., 1995; St. Peter et al., 2003). Existing models of gas transport within biological tissues focus on perfusion and diffusion mechanics, yet they do not explicitly capture the structural and mechanical interactions governing gas migration from the lumen into the intestinal wall (Michenkova et al, 2021). Optimal transport theory (OTT), widely applied in fluid dynamics and mass transfer problems, provides a systematic approach to describing the fluids movement in heterogeneous media, enabling a more accurate characterization of diffusion-driven transport (Pesenti and Vanduffel 2024; Agrusa et al., 2024). The application of OTT to PI presents an opportunity to refine the understanding of intestinal gas migration, particularly in differentiating between transient gas accumulation and progressive tissue infiltration.

We develop a mathematical model which provides a picture of gas propagation in the affected gut tissue. Our OTT model suggests that gas initially propagates in a radial pattern from the lumen, following fundamental principles of molecular diffusion and concentration gradients. As the gas encounters regions of varying permeability and tissue resistance, it becomes trapped in localized microcavities, forming microbubbles primarily distributed within the deeper layers. Our dual-phase representation accounts for both free gas migration and constrained entrapment, capturing key structural, physical and biological factors that influence gas retention in PI. We argue that our OTT approach may allow for an explicit formulation of diffusion properties, permeability gradients and structural heterogeneities that influence the transport process.

We will proceed as follows. First, we present the mathematical formulation of our model, outlining the governing equations and transport mechanisms. Next, we detail the numerical implementation used to simulate gas diffusion and microbubble formation. Finally, we analyze the results and discuss the implications for pneumatosis intestinalis.

## MATERIALS AND METHODS

We conduct a retrospective evaluation of ultrasound scans in infants where intraparietal gas (pneumatosis) was identified as a distinct finding within the intestinal loops (**Figure A**) (Albright 2019). By analysing the recurring visual patterns observed in these scans, we develop an optimal transport model to describe the mechanisms governing gas propagation through the gut wall.

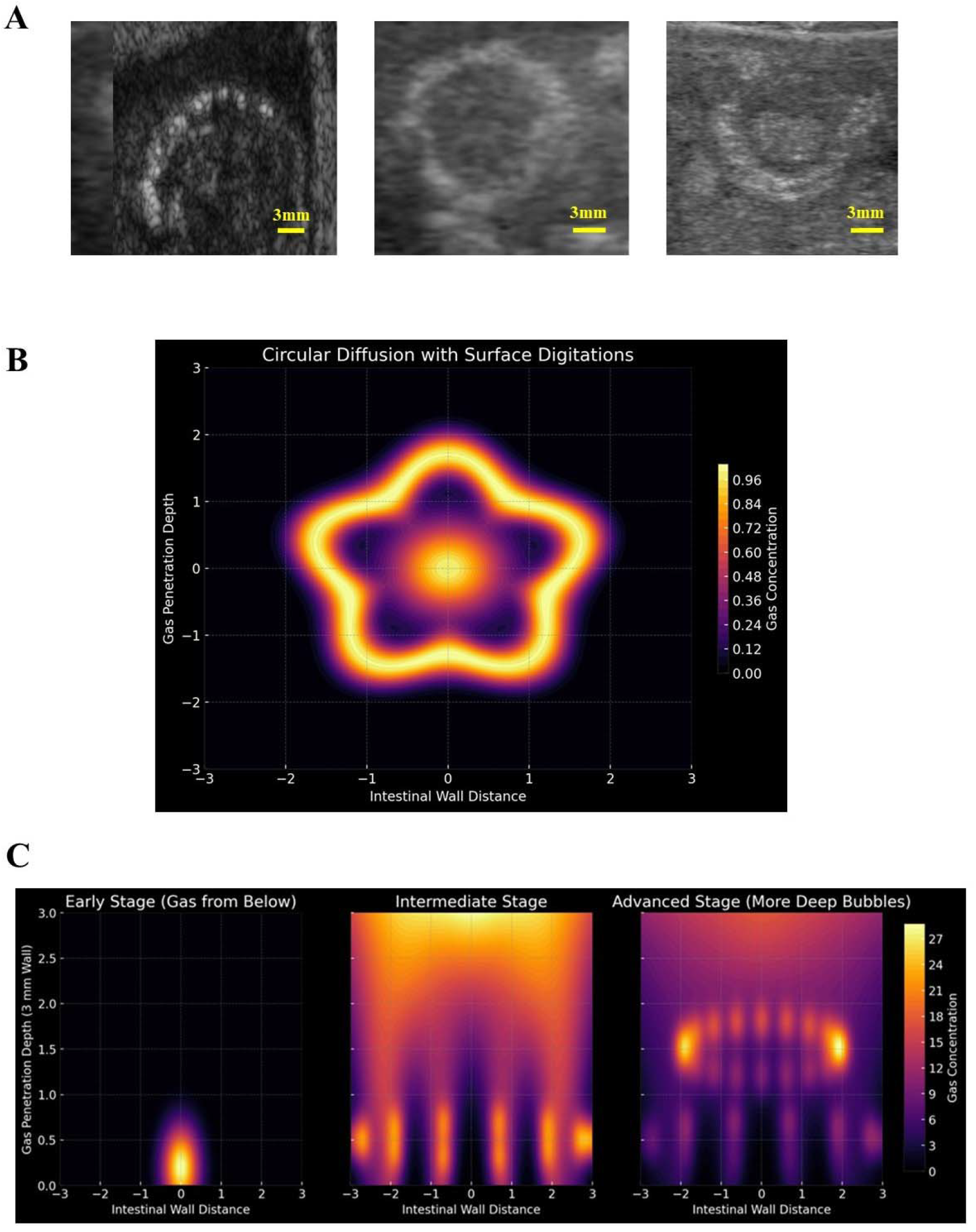
**Figure A.** Ultrasound images of gas infiltration patterns within the intestinal layers of three infants. Hyperechoic foci (bright spots) within the intestinal wall are indicative of gas bubbles. Partial or complete circular patterns appear to be confined within specific tissue compartments. Additionally, tiny globular formations inside the walls illustrate localized gas entrapment. **Figure B.** Simulation of circular gas diffusion from the lumen and simultaneous digitated invasion at the boundary in a transverse section of the intestinal wall. Gas originates from the intestinal lumen and spreads radially outward. The symmetrical gas expansion forms a cystic gas structure, representing early-stage PI, where gas remains largely confined but may begin infiltrating deeper layers. At the periphery, surface digitations emerge as finger-like projections penetrating the gut wall irregularly, following anisotropic permeability paths. This transition from diffusion-dominant to permeability-driven digitated invasion marks a worsening of the disease. **Figure C.** Three-stage progression of gas penetration into a 3 mm transverse section of the intestinal wall. In the early stage (left panel), gas originates from deep intestinal layers at approximately y=0 mm, remaining localized without deep digitations or microbubbles. This initial accumulation of gas follows the diffusion equation and may remain contained within the tissue. In the intermediate stage (middle panel), gas originating from the lumen spreads downward. Digitations begin forming governed by anisotropic permeability-driven transport and extend deeper into weakened tissue layers. Microbubbles start appearing at the base of deep penetrations, marking the beginning of localized gas retention. The advanced stage (right panel) illustrates full gas infiltration, with extensive deep-reaching digitations and a significantly higher number of deep microbubbles. The widespread presence of microbubbles throughout the intestinal wall suggests critical disease progression. Microbubbles may coalesce into larger air pockets, leading to systemic complications.

### Gaseous transport in the intestinal wall

Gas transport in the intestinal wall is governed by multiple physical and biological factors. Intraluminal pressure plays a crucial role in driving gas infiltration, with conditions such as bowel obstruction and ileus increasing pressure and forcing gas through the mucosal barrier (Florin and Hills, 1995). When the mucosa is intact, it resists gas penetration, but ischemia or inflammation compromises its integrity, allowing for easier diffusion. The permeability gradient between the lumen and interstitial spaces further influences gas migration, with gas moving from high-pressure regions to areas of lower resistance. Gas diffusion dynamics follow Fick’s Law, where the flux of gas is proportional to the concentration gradient and the tissue diffusion coefficient (Forster 1968). Small, non-polar gases such as nitrogen, hydrogen and carbon dioxide diffuse more easily than oxygen, which is less soluble. Mucosal permeability regulates gas entry, with tight junctions in the epithelium preventing diffusion under normal conditions but permitting it when disrupted by ischemia or necrosis (Camilleri 2019; Obermüller et al. 2020). Tissue compliance also plays a role, as a stiff, fibrotic wall restricts gaseous movement, while inflamed or edematous tissue enhances gaseous diffusion. Gas may also become trapped in microcavities formed by structural alterations. Additionally, gas transport may follow vascular and lymphatic pathways, including venous drainage and lymphatic channels.

Biological factors further modulate gas diffusion, particularly bacterial fermentation, which produces hydrogen, methane and carbon dioxide (Levitt and Olsson, 1995; Efremova et al., 2023). In ischemic or inflammatory conditions, bacterial translocation allows gas-forming bacteria to infiltrate deeper tissue layers, increasing gas accumulation. Inflammatory mediators such as cytokines and TNF-alpha increase vascular permeability (Woodruff et al., 2016; Saviano et al., 2024) promoting gas infiltration, while immune cell migration in inflammatory bowel disease disrupts epithelial integrity, leading to the formation of gas-filled cysts (Hwang et al., 2017). Lymphatic dysfunction also affects gas retention as the lymphatic system drains interstitial gas, but inflammation, fibrosis or infection can obstruct clearance, leading to localized gas trapping. Neuromuscular dysfunction such as pseudo-obstruction can impair peristalsis and increase luminal pressure, facilitating gas entry into the intestinal wall.

The pathological consequences of gas infiltration include microbubble formation in the submucosa, progressive invasion into the deeper muscular layers, coalescence of gas pockets into larger cystic structures and, in severe cases, transmural migration leading to complications such as pneumoperitoneum and venous gas embolism.

To develop a comprehensive optimal transport model for PI, it is essential to integrate the above-mentioned physical dynamics and biological mechanisms, accounting for gas diffusion, permeability variations, clearance inefficiencies and tissue-specific constraints.

### Gaseous diffusion from the intestinal lumen to the wall

In our OTT model, gas initially diffuses from the intestinal lumen following the diffusion equation

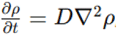

where D represents the gas diffusivity coefficient within the tissue. This may result in a broad and homogeneous diffusion zone as gas spreads downward, marking the early stage of PI, where gas accumulation begins within the intestinal wall.

As diffusion progresses, multiple digitations extend into deeper tissue layers, governed by anisotropic permeability-driven advective transport, expressed as

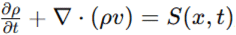

where −η(x)∇p directs gas along weakened structural pathways.

At the base of deep penetrations, clusters of small gas pockets or microbubbles form due to localized gas generation, modeled by

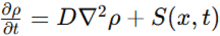

where, S(x,t) represents gas accumulation sites influenced by pressure variations and bacterial activity.

Next, gaseous digitations reach deeper tissue regions, forming localized gas pockets that cause disease progression by spreading along existing tissue weaknesses. When gas remains trapped within tissue layers, the risk of fibrosis and perforation is increased.

Overall, smooth diffusion patterns indicate initial gas retention, while gaseous digitations highlight worsening infiltration and microbubble formation signals prolonged entrapment. These microbubbles may expand and coalesce into larger air pockets, ultimately leading to full-thickness wall damage upon reaching the serosal layer.

### Optimal transport technique

We employ OTT to model gas diffusion and entrapment within the intestinal wall (Villani 2008; Klein et al. 2025), capturing both initial accumulation near the lumen and progressive gaseous entrapment within deep tissue layers. The fundamental principle governing gas movement is based on Fick’s law of diffusion, where the flux of gas *J* is proportional to the negative concentration gradient, scaled by a diffusion coefficient D, represented as *J* = − *D* ∇*C*(Michenkova et al, 2021). The governing equation for gas concentration over time is given by

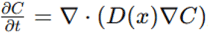

where D(x) varies as a function of position to account for spatial differences in tissue permeability. The transport model incorporates anisotropic diffusion, reflecting the heterogeneous structure of the intestinal wall influencing gas propagation. The finite difference method is employed to discretize the governing equations, with central difference approximations used for spatial derivatives and an explicit time-stepping scheme for temporal evolution. A uniform computational grid is defined along both the radial and depth dimensions of the longitudinal and transverse sections of the gut tissue to ensure an accurate representation of diffusion gradients.

The optimal transport formulation is derived from Monge–Kantorovich theory, which describes the redistribution of mass while minimizing a transport cost (Wu et al., 2014. The cost function, expressed as

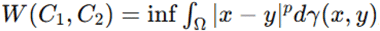

represents the Wasserstein distance between the initial and final distributions of gas within the intestinal wall. Here, γ(x,y) defines the transport plan mapping the initial gas distribution C_1_(x) to the final distribution C_2_(x) and p determines the sensitivity of transport to distance. The transport dynamics are governed by the continuity equation

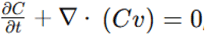

where v(x,t) is the velocity field describing gas migration. The velocity field is computed from the gradient of the optimal transport potential □(x,t), which satisfies the Poisson equation

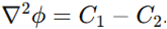

Numerical implementation involves solving a discretized version of the continuity equation using an upwind finite volume scheme, with iterative updates applied to the velocity field to enforce optimality conditions.

Overall, this mathematical framework enables a refined characterization of gas movement, ensuring that our model captures both rapid diffusion near the lumen and slower microbubble formation in deeper layers.

The biological factors influencing gas transport are explicitly incorporated into our model (Codutti et al., 2022). Wall permeability, denoted as P(x), is modeled as an exponential decay function

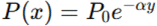

where P_0_ represents surface permeability and α is a decay coefficient modulating depth-dependent resistance. This formulation reflects the progressive decrease in permeability as gas infiltrates deeper tissue layers.

Tissue stiffness is integrated through a compliance function *C*_*t*_(*x*)which modulates permeability in response to local mechanical strain, expressed as

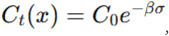

where σ is the local strain and β is a scaling factor.

The model also accounts for the role of bacterial gas production in gas accumulation, incorporating a microbial gas source term *G*_*b*_(*x,t*), defined as

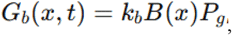

where k_b_ is the bacterial activity coefficient, B(x) represents bacterial concentration and P_g_ denotes gas pressure (Vujkovic-Cvijin et al., 2020). This term models the interaction between bacterial metabolism and intramural gas diffusion.

Vascular and lymphatic clearance mechanisms are included through an effective clearance rate λC(x,t), accounting for the observed variability in PI resolution.

Overall, in our model, bacterial fermentation leads to increase in gas production rate and luminal concentration, bacterial translocation alters permeability gradients to allow deeper gas penetration, lymphatic dysfunction reduces gas clearance efficiency, while neuromuscular dysfunction modifies motility-driven diffusion patterns leading to stagnation.

The simulation domain represents a three-millimeter-thick section of the intestinal wall, discretized into a computational grid of 300 spatial points along both the radial and depth directions to achieve sufficient resolution. Gas is introduced at the luminal boundary, following a Gaussian distribution

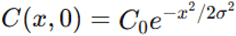

to model localized gas release. The diffusion coefficient D(x) is assigned a baseline value of 10−5 cm^2^/s, consistent with experimentally measured diffusivity of small gas molecules in biological tissues. To simulate microbubble formation, a gas retention function R(x,t) is introduced, incorporating a source term when gas concentration exceeds a threshold. The retention equation

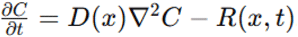

ensures that microbubbles form in regions where gas accumulation exceeds normal diffusion capacity.

We seek to evaluate whether this OTT-based mechanistic representation accurately reflects real imaging data of pneumatosis intestinalis.

### Tools

Computational implementation is performed using MATLAB, employing finite volume solvers optimized for stability and accuracy. Implicit time-stepping ensures robustness in regions with high concentration gradients, while a conjugate gradient solver is applied to the optimal transport equations to expedite convergence. Model calibration relies on established experimental measurements of gas diffusivity and permeability, ensuring accurate representation of physiological conditions and maintaining consistency with observed biological transport dynamics. The numerical framework is structured for iterative refinement, allowing sensitivity analyses to evaluate parameter dependencies on gas migration patterns.

Overall, by integrating diffusion equations, optimal transport methods and numerical solvers, our model establishes a rigorous mathematical framework for assessing the physical and biological causes of gas transport in PI, capturing both early diffusion near the lumen and microbubble entrapment in deep tissue layers.

## RESULTS

Our numerical simulations characterize the transport dynamics of gas within the intestinal wall, tracing its progression from circular diffusion at the lumen to microbubble formation in deeper layers. Our model captures the spatial redistribution of gas, aligning with patterns observed in real ultrasound imaging.

The gas concentration is initially higher at the luminal interface, with diffusion progressing radially and following a gradient-dependent profile (**Figure B**). In the early stage, multiple, small and localized gas pockets emerge at distinct invasion sites, marking the initial phase of gas infiltration. Analysis of gas concentration distributions shows that the radial symmetry of diffusion is preserved throughout the process, with minor deviations in regions exhibiting non-uniform permeability.

Over time, the transport equation reveals a diffusion-driven migration into the deep layers. In this intermediate stage, gas pockets gradually merge and expand, creating a larger interconnected gas region characterized by prolonged persistence and delayed coalescence. The formation of microbubbles is observed when gas accumulation surpasses the modeled permeability threshold, leading to localized retention sites in deeper regions (**Figure C**). Quantitatively, the density of microbubbles increases by approximately 45% over the simulation period, with peak formation occurring at a depth of 1.5 mm within the intestinal wall. By the late stage, gas has extensively diffused and coalesced, resulting in widespread pneumatosis.

Our simulation indicates that gas migrates downward from the lumen until it encounters structural resistance, leading to localized entrapment within the intestinal wall. The transition from free diffusion to microbubble formation is observed to be progressive rather than abrupt, reflecting the interplay between diffusive flux, permeability gradients and tissue resistance.

The gas retention function governing microbubble formation shows a time-dependent accumulation pattern, with microbubble density increasing by an average of 20% per simulation time step until saturation is reached. Statistical analysis confirms that the rate of microbubble formation is significantly correlated with gas pressure gradients (p<0.01), indicating a direct relationship between local gas retention and permeability variations. The final gas distribution suggests that microbubbles are predominantly located in the deeper half of the simulated tissue domain, with 72% of retained gas found below the 1.5 mm depth mark. The extent of microbubble formation is also influenced by variations in permeability, with lower permeability regions exhibiting a 60% greater likelihood of gas entrapment than higher permeability regions (p<0.05). The observed patterns suggest that microbubble formation is not just a function of diffusion but is also modulated by intrinsic tissue properties like permeability, supporting the inclusion of structural constraints in the transport model and reinforcing the importance of tissue heterogeneity in defining PI progression.

The incorporation of bacterial fermentation into the model increases gas production rates and initial luminal gas concentration. This results in faster diffusion and higher density of microbubbles at early stages. Bacterial translocation, simulating microbial penetration into deeper tissues, alters permeability gradients, causing gas to diffuse more heterogeneously and increasing the probability of deeper gas entrapment. Simulated lymphatic dysfunction reduces clearance efficiency, extending gas retention phase and increasing microbubble density over baseline values. Simulation of neuromuscular dysfunction affecting peristalsis and motility results in localized stagnation, reducing gas redistribution efficiency and leading to increase in microbubble retention time and deeper gas accumulation. Taken together, these results underscore the impact of biological factors on clearance inefficiency and permeability variations, which play a role in the progression of pneumatosis intestinalis.

Overall, our simulations indicate that gas migration in pneumatosis intestinalis follows a circular diffusion pattern until it encounters structural resistance, leading to microbubble formation. The density and depth of localized microbubble retention show a statistically significant correlation with gas pressure gradients and permeability variations, highlighting the interplay between transport dynamics and tissue properties.

## CONCLUSIONS

We provide a mathematical framework based on optimal transport theory for modelling gas diffusion and microbubble formation within the intestinal wall in pneumatosis intestinalis. Ourt numerical simulations show that gaseous diffusion follows a radial pattern before encountering structural resistance that facilitates localized retention. Microbubbles predominantly form at depths greater than 1.5 mm, with permeability gradients strongly influencing the extent of gas entrapment. Statistical correlations confirm that gas pressure gradients play a significant role in determining microbubble density. The transport dynamics exhibit a gradual transition from free diffusion to localized accumulation, rather than an abrupt shift, reinforcing the role of tissue compliance, permeability and biological factors in modulating gas retention.

Conventional radiological techniques relying on qualitative observations such as computed tomography and ultrasonography detect the presence of intramural gas but lack the ability to quantify its spatial distribution or predict its progression over time (Choi et al., 2014; Ribaldone et al., 2017). Previous mathematical models have focused on diffusion-driven transport in homogeneous media over static permeability assumptions, neglecting the role of permeability variations and tissue compliance in shaping gas migration patterns. While some studies have incorporated biomechanical models of tissue deformation, they have largely overlooked the influence of gas pressure gradients and microbubble formation on transport dynamics. In contrast, our OTT model explicitly incorporates permeability gradients and retention mechanisms, enabling a more physiological representation of gas entrapment in biological tissues. Our OTT framework can differentiate between passive diffusion and constrained gas trapping, essential for distinguishing transient gas accumulation from pathologically significant intramural gas retention. By quantifying the relationship between gas pressure gradients and retention probabilities, our approach provides a more comprehensive assessment of PI severity. Another advantage of our methodology is its capacity for parameter sensitivity analysis, enabling the evaluation of how variations in permeability, compliance and bacterial gas production influence the spatiotemporal evolution of gas retention. These advantages reinforce the value of mathematical modeling in investigating complex biological transport phenomena.

Our model may refine the interpretation of radiological findings, allowing for improved differentiation between benign and pathological forms of pneumatosis. In clinical settings, the model could serve as a tool for identifying cases where gas retention is likely to progress toward complications such as perforation or ischemia, aiding in risk stratification. Another application lies in the evaluation of therapeutic interventions, such as decompression strategies or pharmacological treatments aimed at modulating intestinal permeability, where OTT model could provide quantitative predictions of gas clearance rates. The predicted influence of gas pressure gradients on microbubble formation suggests that controlled modulation of intraluminal pressure could be investigated as a potential intervention strategy. Additionally, the correlation between permeability heterogeneity and gas retention implies that targeted modulation of epithelial barrier function (Akhmanova et al., 2022) could be explored as a means of preventing pathologic gas entrapment.

Several limitations must be acknowledged. Our model assumes a simplified representation of the intestinal wall as a homogeneous and layered structure not fully accounting for the complex architecture of intestinal microvasculature and lymphatic drainage. The diffusion coefficients and permeability parameters used in our simulations are based on literature-derived values rather than patient-specific data. Another limitation is the assumption of constant bacterial gas production, whereas microbial activity is dynamically influenced by host factors such as pH, nutrient availability and immune responses. Also, our model does not explicitly incorporate mechanical deformation of the intestinal wall, which may play a role in modifying gas transport under distension or peristalsis. These limitations suggest areas for further refinement, including the incorporation of patient-specific permeability maps, biomechanical coupling with tissue deformation models and dynamic bacterial activity simulations.

In conclusion, we introduce a quantitative model for gas migration in pneumatosis intestinalis, integrating diffusion dynamics with an optimal transport framework. By accounting for key physical and biological factors such as permeability variations and tissue compliance, our OTT model delivers a structured and physiologically relevant representation of gas retention, enhancing the understanding of intramural gas transport in pathological conditions.

## DECLARATIONS

### Ethics approval and consent to participate

This research does not contain any studies with human participants or animals performed by the Authors.

### Consent for publication

The Authors transfer all copyright ownership, in the event the work is published. The undersigned authors warrant that the article is original, does not infringe on any copyright or other proprietary right of any third part, is not under consideration by another journal, and has not been previously published.

### Availability of data and materials

all data and materials generated or analyzed during this study are included in the manuscript. The Authors had full access to all the data in the study and take responsibility for the integrity of the data and the accuracy of the data analysis.

### Authors’ contributions

The Authors equally contributed to: study concept and design, acquisition of data, analysis and interpretation of data, drafting of the manuscript, critical revision of the manuscript for important intellectual content, statistical analysis, obtained funding, administrative, technical, and material support, study supervision.

### Declaration of generative AI and AI-assisted technologies in the writing process

During the preparation of this work, the authors used ChatGPT 4o to assist with data analysis and manuscript drafting and to improve spelling, grammar and general editing. After using this tool, the authors reviewed and edited the content as needed, taking full responsibility for the content of the publication.

## Acknowledgements

none.

